# Draft Genome Sequence and intraspecific diversification of the wild crop relative *Brassica cretica* Lam. using demographic model selection

**DOI:** 10.1101/521138

**Authors:** Antonis Kioukis, Vassiliki A. Michalopoulou, Laura Briers, Stergios Pirintsos, David J. Studholme, Pavlos Pavlidis, Panagiotis F. Sarris

## Abstract

Crop wild relatives contain great levels of genetic diversity, representing an invaluable resource for crop improvement. Many of their traits have the potential to help crops become more resistant and resilient, and adapt to the new conditions that they will experience due to climate change. An impressive global effort occurs for the conservation of various wild crop relatives and facilitates their use in crop breeding for food security.

The genus *Brassica* is listed in Annex I of the International Treaty on Plant Genetic Resources for Food and Agriculture. *Brassica oleracea* (or wild cabbage) is a species native to coastal southern and western Europe that has become established as an important human food crop plant because of its large reserves stored over the winter in its leaves.

*Brassica cretica* Lam. is a wild relative crop in the brassica group and *B. cretica* subsp. *nivea* has been suggested as a separate subspecies. The species *B. cretica* has been proposed as a potential gene donor to a number of crops in the brassica group, including broccoli, Brussels sprout, cabbage, cauliflower, kale, swede, turnip and oilseed rape.

Here, we present the draft *de novo* genome assemblies of four *B. cretica* individuals, including two *B. cretica* subsp. *nivea* and two *B. cretica*.

*De novo* assembly of Illumina MiSeq genomic shotgun sequencing data yielded 243,461 contigs totalling 412.5 Mb in length, corresponding to 122 % of the estimated genome size of *B. cretica* (339 Mb). According to synteny mapping and phylogenetic analysis of conserved genes, *B. cretica* genome based on our sequence data reveals approximately 30.360 proteins.

Furthermore, our demographic analysis based on whole genome data, suggests that distinct populations of *B. cretica* are not isolated. Our findings suggest that the classification of the *B. cretica* in distinct subspecies is not supported from the genome sequence data we analyzed.

## Introduction

### Crop Wild relatives

Many plant species are used in the food and agriculture market; however, 30 crops account for the 95% of food production worldwide (Brozynska *et al.* 2016). Domesticated crops, used in the food production, show reduction in the genetic diversity, compared to their respective <u>C</u>rop <u>W</u>ild <u>R</u>elatives (CWRs). This genetic “bottleneck” of domestication (Tanksley and McCouch 1997) resulted to loss of valuable genetic alleles. On the other hand, during the domestication process cultivated varieties, introgression from wild species may generate additional genetic diversity may arise (Hufford *et al.* 2013, Sawler *et al.* 2013).

As wild ‘progenitors’ of crops continue to evolve under abiotic and biotic stresses, it is very important to conserve this resulting genetic biodiversity, which can be useful for agriculture (*in situ* conservation). Seed banks or germplasm collections are also important to preserve as another resource for agriculture (*ex situ* conservation). The total genome sequencing of CWRs may be used first to characterize wild populations and inform strategy for their conservation. On the other hand, analysis of the sequence can reveal genetic variation and important genetic characters that have been lost during domestication, and that could be transfer into crop species to support food security, climate adaptation and nutritional improvement (Brozynska *et al.* 2016). The ready availability of low-cost and high-throughput re-sequencing technologies enable the survey of CWR genomes for genetic variation and novel genes and allelels. Recent decades have seen some remarkable examples of introducing favored traits from CWRs into their respective domesticated crop plants. In most cases, these traits concern resistance to biotic stresses, such as resistance to late blight (*Phytophthora infestans)* from the wild potato *Solanum demissum* Lindl. (Prescott-Allen 1986, Witek *et al.* 2016). Besides biotic tolerance, many quantitative trait loci have been identified and/or introduced, regarding the grain quality for increased yield, such as from *Oryza rufipogon*, a wild species of rice, to *Oryza sativa* (Septiningsih *et al.* 2003) and grain hardness from *Hordeum spontaneum* (wild barley) (Li *et al.* 2010).

### Brassica oleracea: Crops and Genomic features

*Brassica oleracea* L. is a very important domesticated plant species, comprising of many vegetable crops as different cultivars, such as cauliflower, broccoli, cabbages, kale, Brussels sprouts, savoy, kohlrabi and gai lan. *Brassica oleracea* or wild cabbage belongs to the family of *Brassicaceae* and is found in coastal Southern and Western Europe. The species has become very popular because of its high content to nutrients, such as vitamin C, its anticancer properties (Higdon *et al.* 2007), as well as the high food reserves in its leaves.

*Brassica oleracea* constitutes the one of the three diploid *Brassica* species in the classical triangle of U (1935) (genome: CC), that contains nine chromosomes. The other two species in this group are *B. rapa* (L.) L. (genome: AA) with 10 chromosomes and *B. nigra* (L.) W. D. J. Koch (the black mustard) (genome: BB) with 8 chromosomes.

These three species, as they are closely related, gave rise to new allotetraploids species that are very important oilseed crops, the *B. juncea* (genome: AABB), *B. napus* L. (genome: AACC) and *B. carinata* (genome: BBCC). There is evidence for each of the *Brassic*a genomes to have undergone a whole-genome duplication (Bowers *et al.* 2003, Jiao *et al.* 2011) and a *Brassiceae*-lineage-specific-whole-genome triplication, which was followed after the divergence from *Arabidopsis* lineage (Lysak *et al.* 2005, Wang *et al.* 2011).

In 2014, Liu *et al*., reported a draft genome of *B. oleracea* var. *capitata* and a genomic comparison with its very close sister species *B. rapa*. A total of 45,758 protein-coding genes were predicted, with mean transcript length of 1,761 bp and 3,756 non-coding RNAs (miRNA, tRNA, rRNA and snRNA). It is observed that there is a greater number of transposable elements (TEs) in *B. oleracea* than in *B. rapa* as a consequence of continuous amplification over the last 4 million years (MY), the time that the two species were diverged from a common ancestor, whereas in *B. rapa* the amplification is made mostly in the recent 0.2 MY (Liu *et al.* 2014). Moreover, there has been observed massive gene loss and frequent reshuffling of triplicated genomic blocks, which favored over-retention of genes for metabolic pathways.

### Brassica cretica

Among the Aegean islands, Crete is the largest and the most diverse from a floristically point of view. It has experienced a much longer history of isolation compared to the smaller Aegean islands. Over two-thirds of all Greek plant species are found in Crete and it has the greatest proportion of endemic species in the Aegean area (Edh *et al.* 2007, Greuter 1971, Webb 1978). Crete was separated from the mainland of Greece around 8 million years ago (Edh *et al.* 2007; Creutzburg 1963, Dermitzakis & Papanikolaou 1981). For many Cretan plant species suitable habitat is restricted at present to areas that are surrounded by a ‘sea’ of low-lying areas acting as dispersal barriers (Davis 1951), this especially includes various chasmophytic plant species. *Brassica cretica* Lam is a typical example of a Cretan chasmophyte species. It is a wild plant species preferentially inhabiting limestone cliffs and gorges, mainly in Crete but also on other Mediterranean countries in the surrounding coastal areas (Snogerup *et al.* 1990). *Brassica cretica* Lam, is wild relative of the cultivated cabbage, *B. oleracea* L. (Lázaro and Aguinagalde, 1998). The species is hermaphrodite (has both male and female organs) and is pollinated by Insects. *Brassica cretica* is a diploid (2n = 18), partially self-incompatible, with a native distribution in Greece (mainly Crete and North Peloponnisos). The plants are perennial and up to 150 cm high, with white or yellow, insect-pollinated flowers that develop into siliqua. Preliminary analyses of electrophoretic variation show that *B. cretica* is outcrossing (few deviations from Hardy-Weinberg equilibrium) and that populations on Crete have undergone extensive divergence at allozyme loci (Lázaro and Aguinagalde, 1998). The geographical isolation has been proposed as the main reason of the significant differences observed among the local *B. cretica* populations for a number of morphological traits (Snogerup *et al.* 1990, Widén *et al.* 2002). Furthermore, flower colour differences could constitute an additional mechanism of genetic isolation among populations if different pollinators prefer different types of flower (cf. Grant 1994). However, the rates of migration among *B. cretica* populations have not been properly quantified, which makes unclear whether the low gene flow alone could explain the population divergence, or if the local adaptation (divergent selection) must be invoked. Widen and colleagues, (Widén *et al.* 2002) reported that the high levels of differentiation observed at allozyme loci and quantitative traits among Cretan *B. cretica* populations, were in agreement with the idea of non-adaptive differentiation combined with limited gene flow. However, the allozymes may not provide accurate assessments of population structure and gene flow, since, evidence has been reported that at least one allozyme locus is under divergent selection in a variety of species (Pogson *et al.* 1995, Riginos *et al.* 2002, Dhuyvetter *et al.* 2004, Edh *et al.* 2007). Moreover, Edh *et al.,* (2007), using nuclear and chloroplast microsatellite markers, studied the differentiation of seven Cretan populations of *B. cretica* and concluded that current patterns of diversification in *B. cretica* are mainly a result of genetic drift.

*Brassica cretica* Lam. is being considered as a wild crop relative of a big number of crops of the genus *Brassica*, proposed to be the ancestor of broccoli, Brussel sprouts, cabbage, cauliflower, kale, swede, turnip and oilseed rape. Since this species is thought to be the gene donor of many crops of the brassica group, it might contain genes that are not included in the domesticated crops, as well as, a different set of NLRs. Potential analysis of the NLRsome of wild species would help us find which genes or locus are responsible for the recognition of effectors from important phytopathogens and thus create resistant plants in the field via transfer of these favored genes/locus (Chen *et al.* 2013).

### Aim of this work

Here, we present the first draft de novo genome assemblies of four individual of *B. cretica* and using the derived genomic data we investigate mechanisms of diversification of four isolated *B. cretica* populations taking into consideration their genomic and subspecies variation.

## Materials and Methods

### Plant Material

Due to the high phenotypic variability of *B. cretica*, a number of subspecies and varieties have been defined. Snogerup *et al*., (1990) recognize three subspecies of *B. cretica*: subsp. *aegea*, subsp. *cretica*, and subsp. *laconica*, whereas Gustafsson *et al*., (1976) suggest only two subspecies, subsp. *cretica* and subsp. *nivea* (sometimes referred to as *B. cretica* subsp. *cretica* var. *nivea*; Turland *et al.* 1993), which includes (pale) yellow and white-flowered variants, respectively.

According to the Vascular Flora of Greece (Dimopoulos *et al.* 2013) there are three subspecies *B. cretica* subsp. *aegaea* (Heldr. & Halácsy; Snogerup; Gust & Bothmer), *B. cretica* subsp. *cretica* and *B. cretica* subsp. subsp. *laconica* (Gust. & Snogerup), while *B. cretica* subsp. *nivea* (Boiss & Spruner; Gust. & Snogerup) and *B. nivea* (Boiss & Spruner) are considered as synonym and misapplied of *B. cretica* Lam. subsp. *cretica*, which has been reported for the mainland of Greece and for the floristic region of Crete and Karpathos (Dimopoulos *et al.* 2013).

For the present study three mainland and one island population of *B. cretica* from Greece have been studied. Two *B. cretica* subsp. *nivea* (Boiss & Spruner) M. A. Gust. & Snogerup individuals from the first two mainland populations respectively (A, B) and two *B. cretica* Lam. individuals, one from the third mainland population (C) and the other from Crete, the island population (D), have been used for the genome assembly (Figure 1).

**Figure 1:**
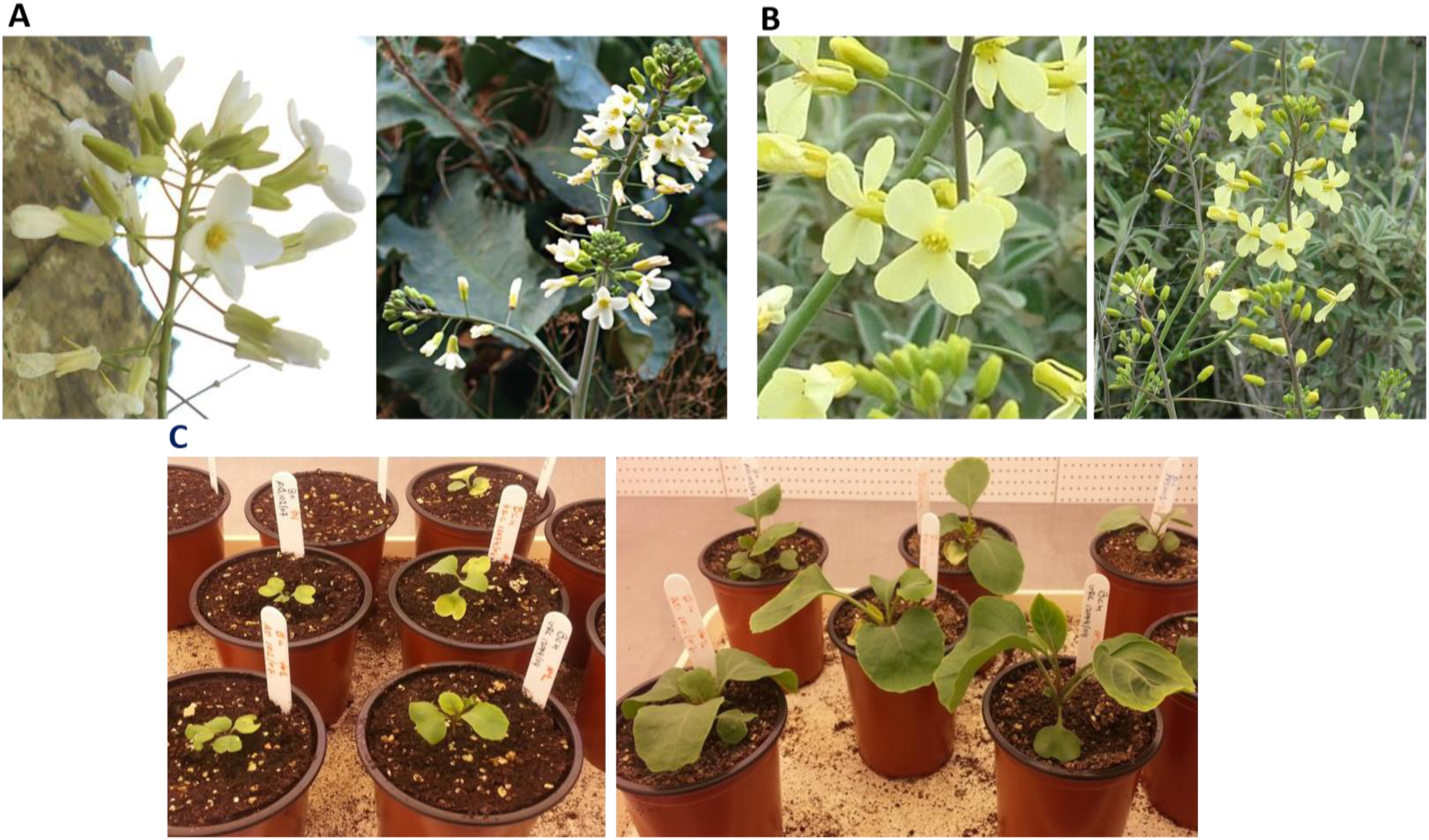
**A:** *Brassica cretica* subsp. *nivea*; **B:** *Brassica cretica*; **C:** *Brassica cretica* in the plants’ growth chamber.

### Total DNA extraction and Library preparation for NGS

Genomic DNA was extracted from the young emerging leaves using two previously published protocols. For total DNA isolation of up to 1 g plant leaf tissue was used. For the DNA isolation we used several protocols including the DNeasy Plant Mini Kit from Qiagen, as the manufactures propose. Likewise, we used a modified triple CetylTrimethyl Ammonium Bromide (CTAB) extraction protocol for total plant DNA isolation, as it has been described before (Abbasi and Afsharzadeh 2016).

The yield and quality of DNA were assessed by agarose gel electrophoresis and by a NanoDrop spectrophotometer (NanoDrop Technologies, Wilmington, Delaware) and quantified by Qubit broad range assay (Thermo Fisher Scientific). Illumina sequencing libraries were prepared, after fragmenting 500 ng of DNA to an average size of 500 bp, using Nextflex8 Rapid DNAseq kit for Illumina sequencing (Bioo Scientific) with adapters containing indexes and 5–8 cycles polymerase chain reaction (PCR) (Head *et al.* 2014). Library quality was determined using D1000 screen-tapes (Agilent) and libraries were either sequenced individually or combined in equimolar pools.

### Genome assembly and annotation

Prior to assembly, Illumina MiSeq sequence reads were filtered on quality scores and trimmed to remove adapter sequences using Trim Galore (https://www.bioinformatics.babraham.ac.uk/projects/trim_galore/) with q = 30 (Quality Phred score cutoff = 30). Reads were assembled into contigs using SOAPdenovo2 (Luo *et al.* 2012) with k = 127 (k-mer value = 127). Configuration files used for the SOAPdenovo2 assembly can be found on FigShare at under DOI 10.6084/m9.figshare.7583396. Contigs shorter than 500 bp in length were removed.

Protein-coding genes were predicted in the draft genome of individual A using the MAKER pipeline version 2.31.10 (Cantarel *et al.* 2008), incorporating ab initio prediction with Augustus 3.1 (Stanke and Waack 2003). Configuration files for the MAKER annotation can be found on FigShare under DOI 10.6084/m9.figshare.7583672. The GFF file generated by MAKER was converted into NCBI’s Feature Table (.tbl) format using Genome Annotation Generator (Geib *et al.* 2018) version 2.0.1.

For comparison with *B. oleracea* var. *oleracea* (wild cabbage) and for variant calling between *B. cretica* individuals, we aligned *B. cretica* MiSeq reads against the previously published reference genome sequence (GenBank: GCA_000695525.1) (Parkin *et al.* 2014) using BWA (Li and Durbin 2009). SNP calling was performed as previously described (Yemataw *et al.* 2018). That is, we identified single-nucleotide polymorphisms by alignment against the reference genome sequence, according to the following procedure. After trimming and filtering with TrimGalore, sequence reads were aligned against the reference sequence using Burrows-Wheeler Aligner (BWA) (Li and Durbin 2009) mem version 0.7.15-r1140 with default options and parameter values. Candidate SNVs were identified using Sequence Alignment/Map tools (SAMtools)/binary call format tools (BCFtools) package, version 1.6 (Li *et al.* 2009), using the following command-lines:

`samtools mpileup -u -f genome.fasta alignment.bam 4 alignment.bcf` and `bcftools call -m -v –Ov alignment.bcf 4 alignment.vcf`

The candidate variants were then filtered using the following command line: `bcftools filter –SnpGap 100 –include ‘(REF=“A” | REF=“C” | REF=“G” | REF=“T”) & %QUAL >35 & MIN(IDV)>2 & MIN(DP)>5 & INDEL=0’ alignment.vcf 4 alignment.filtered.vcf`

This filtering step eliminates indels with low-confidence single-nucleotide variant calls. It also eliminates candidate SNVs within 10 base pairs of an indel, since alignment artefacts are relatively common in the close vicinity of indels. Allele frequencies at each SNP site were estimated from frequencies of each base (adenine (A), cytosine (C), guanine (G) or thymine (T)) among the aligned reads. Thus, we would expect an allele frequency of close to zero or one for homozygous sites and approximately 0.5 for heterozygous sites in a diploid genome. The binary alignment/map (BAM)-formatted BWA-mem alignments were converted to pileup format using the samtools mpileup command in SAMtools version 1.6 (Li *et al.* 2009), with default options and parameter values. From the resulting pileup files, we used SNPsFromPileups version 0.1 (https://github.com/davidjstudholme/SNPsFromPileups) to detect SNPs. For SNP detection, we considered only sites where depth of coverage by aligned reads was at least 5× for all four datasets.

### Genome annotation

Genome annotation was performed using the MAKER pipeline (Campbell *et al.* 2014, Cantarel *et al.* 2008) version 2.31.10. Ab initio gene prediction was performed using Augustus (Stanke and Waack 2003) version 3.1 trained on *Arabidopsis*. Amino acid sequences predicted by MAKER were subjected to analysis with PfamScan to identify those predicted proteins containing an NB-ARC domain (Finn *et al.* 2014).

### Allele Frequency Spectrum (AFS)

The AFS defined as ξ = {ξ_i_: number of sites with derived allele counts being *i*} is a useful summary of the data especially for demography inference. To calculate the AFS, we mapped the reads of *B. cretica* to the *B. oleracea* reference genome. This allowed us to use all specimens and also to use the *B. oleracea* as an outgroup that denotes the ancestral state. Following the GATK best practices pipeline (Van der Auwera *et al.* 2013), this mapping resulted in approximately six million single nucleotide polymorphisms (SNPs).

*Brassica oleracea* has been examined thoroughly in the past and there is a gene list of the organism organized into chromosomes. We used this list to exclude SNPs with a distance less than 10kb from those coding regions. This process of removing SNPs is necessary when the SNPs are used to infer the demographic model. Due to linkage disequilibrium SNPs within or in the proximity of genic regions are affected by selection forces, especially negative selection. Negative selection effectively increases the low frequency derived variants and therefore it introduces biases in the demographic inference. For this reason, we excluded SNPs located within or in the proximity of genic regions.

### Demographic inference

Using *dadi* to infer the demographic model.

Inferring a demographic model consistent with a particular data set requires random walks into a large parameter space by simulating the model using Monte Carlo coalescent-theory based approaches. The most well-known approach based on Monte Carlo coalescent simulations is the Approximate Bayesian Computation (ABC) inference (Beaumont 2002). The main handicap of these methods is their scalability to genome-wide size data sets. Another issue arises when multiple populations are free to interact through migration (either symmetric or asymmetric) resulting in an increase of the parameters and, therefore, the required complex calculations. These complexities hinder any effort to thorough explain the statistical properties of the summary statistics produced during the walks. To avoid these problems we based our demographic model inference on the multi-population allele frequency spectrum (AFS) (Bustamante *et al.* 2001, Caicedo *et al.* 2007, Hernandez *et al.* 2007, Nielsen *et al.* 2009) due to the fact that demographic history of a population is reflected in the allele frequency spectrum. By comparing the different spectra produced by simulations and observations we can access the model’s goodness of fit and estimate the best parameter values for each model.

In spite of the existence of efficient algorithms for the simulation of a single population AFS (Adams and Hudson 2004, Marth *et al.* 2004, Williamson *et al.* 2005), the joint AFS between two or more populations still requires very computationally intensive coalescent simulations. For more than two populations the computational complexity becomes prohibitively large. Approximations of the joint-AFS using a numerical solution of a diffusion equation have been used extensively in the past (Der Sarkissian *et al.* 2015), enabling simulations of a joint-AFS for two populations in a reasonable computation time. Although the diffusion approach neglects linkage disequilibria, we can use composite likelihood function as a consistent estimator for evaluating genetic scenarios. Concerns about the use of composite likelihood in population genetics are overcome by allowing conventional and parametric bootstrap of the data.

The *dadi* python package (Gutenkunst *et al.* 2009) implements these approximations and in conjunction with the *dadi_pipeline* described in (Portik *et al.* 2017) allows for adequate exploration of the parameter space. The *dadi_pipeline* consists of three optimization rounds and a final plotting step. We used 30 demography models ranging from simple (populations never diverge) to complex (ancient divergence with asymmetric migrations between the two populations) to find the best fitting model. These demographic models comprise a thorough list of two population possible models and they were first examined by Portik *et al.* 2017).

The initial two rounds of optimizations search the parameter space for the parameter set that best describes the data under each of the thirty models. For every model we sampled 50 different parameter sets and 50 repetitions of the each set to get the actual global maximum for each model while avoiding local maxima. We based our selections of the best parameter values on the AIC score for each model. To assess which demographic model better reflects the true demographic history of the *B. cretica* population a simple comparison between the respective AIC scores from each model is not valid because AIC is not comparable between non-nested models. We compared the models using Akaike weights (Wagenmakers *et al.* 2004), by calculating the difference between each model’s AIC and the AIC of the best candidate model. With a simple transformation we can calculate an estimate of the relative likelihood L_i_ of each model *i* and by dividing each Li with the sum of Li we can normalize the weights and compare the models, and therefore we can find the model that better fits the data (Wagenmakers *et al.* 2004).

### GenBank genomes assembling deposition details

The assembly statistics for each of the assembled genomes can be found in Table 1 and Table S4. The assembly accession number as they appear at the GenBank are: 1) *Brassica cretica* PFS-1207/04, GenBank assembly accession: **GCA_003260655.1**; 2) *Brassica cretica* PFS-001/15, GenBank assembly accession: **GCA_003260635.1**; 3) *Brassica cretica* PFS-109/04, GenBank assembly accession: **GCA_003260675.1**; 4) *Brassica cretica* PFS-102/07, GenBank assembly accession: **GCA_003260695.1**.

**Table 1:** Assembly Statistics.

|  | *Bc* PFS-1207/04 | *Bc* PFS-001/15 | *Bc* PFS-109/04 | *Bc* PFS-102/07 |
| --- | --- | --- | --- | --- |
| **Total sequence length - bp** | 412,521,210 | 208,353,552 | 434,935,090 | 400,211,799 |
| **Number of contigs** | 243,461 | 100,644 | 338,759 | 396,633 |
| **Contig N<sub>50</sub> - bp** | 2,820 | 2,197 | 1,572 | 1,027 |
| **Contig L<sub>50</sub> - bp** | 36,638 | 25,402 | 57,226 | 100,090 |
| **Number of contigs** | 243,461 | 100,644 | 338,759 | 396,633 |
| **Genome coverage depth** | 55.0x | 63.0x | 64.0x | 39.0x |

## Results and Discussion

### Demographic Model Inference

Demographic analysis based on whole genome data suggests that populations of *B. cretica* are not isolated. In agreement with others, we suggest that the classification of the *B. cretica* in distinct subspecies is not supported from the data. Using only the non-coding part of the data (thus, the part of the genome that has evolved nearly neutrally), we find the gene flow between different *B. cretica* population is rather recent and its genomic diversity is high.

We followed two approaches, to infer the neutral demographic model for the *B. cretica* data. The two approaches are related to the separation of the individual plants into distinct groups (i.e., population or subspecies). According to the first approach, the subspecies approach, we separate the individuals into two groups specified by their subspecies definition. Plants A and B are characterized as *B. cretica* subsp. nivea SFP1207/94 and *Brassica cretica* subsp. nivea SFP0001/15 (Cretan isolate), respectively, and they constitute group 1, whereas plants C and D are *B. cretica* SFP109/07 and *B. cretica* SFP102/07, respectively, and they define group 2. The second approach is based on the PCA plot of the data, which depends on the differences at the DNA level. We call the second approach the genetic approach. We applied logistic Principal Component Analysis (http://arxiv.org/abs/1510.06112v1) (logPCA) since the polymorphisms at each site define a binary state. The results of the logPCA are shown in Figure 2.

**Figure 2:**
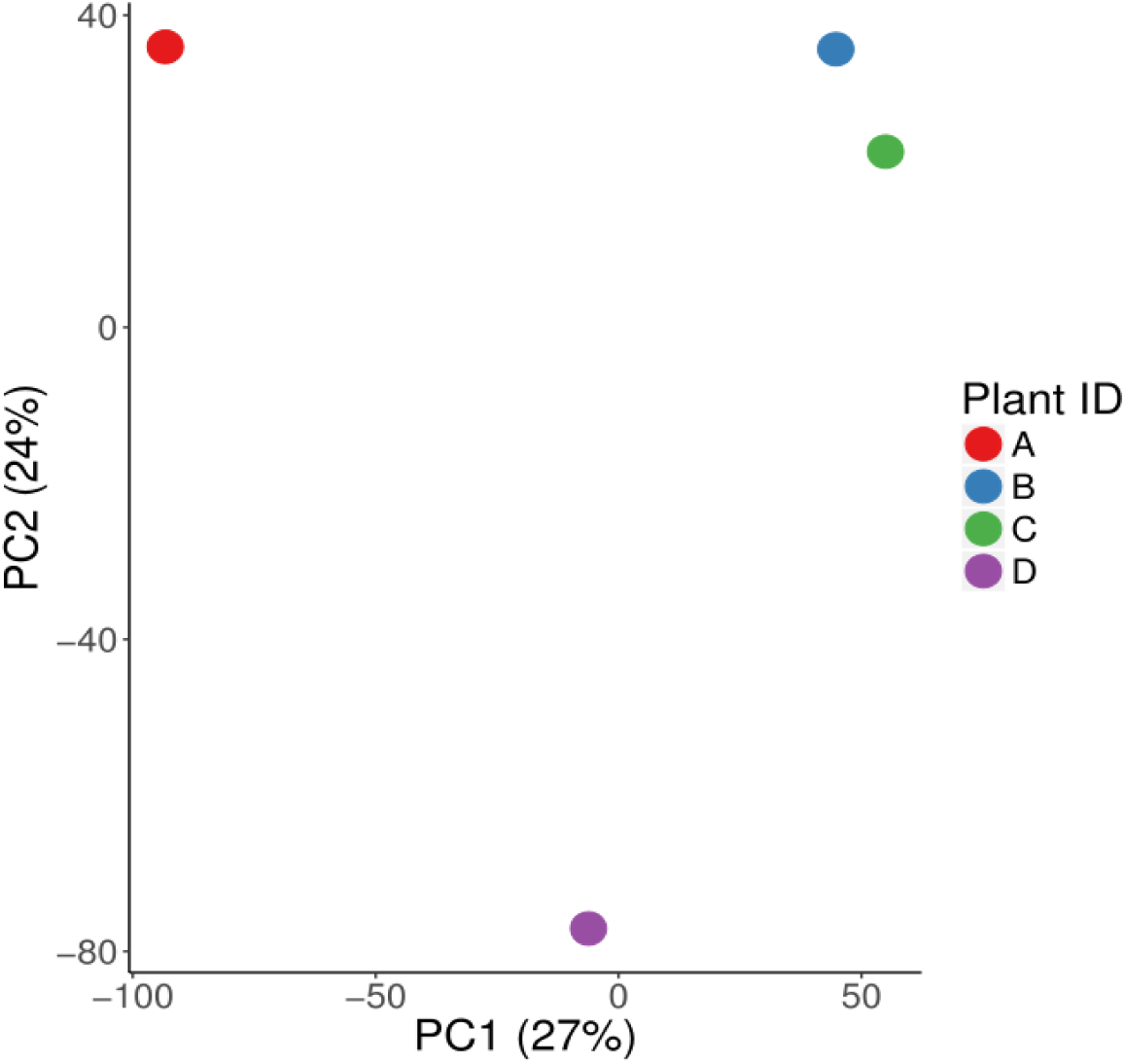
The logPCA results of binary SNP data at the level of the first two Axes.

Along the PC1 we defined the members of 2 populations. Population 1 consists of plant A, whereas population 2 of plants B, C, and D. The PC1 and PC2 explain 51% of the data variance.

### Demographic Model Inference based on the subspecies definition

Following the subspecies definition of the two groups of plants, the model “Vicariance with late discrete admixture” is the most likely among the 30 different models with two populations. Such a model suggests that the two subspecies were discrete for a long period of time. However, recently, introgression took place from group 1 (plants A and B) to group 2. Such a massive gene flow suggests that the two groups of plants may not define distinct subspecies, therefore they can be considered as different population of the same species (Figure 3A).

**Figure 3:**
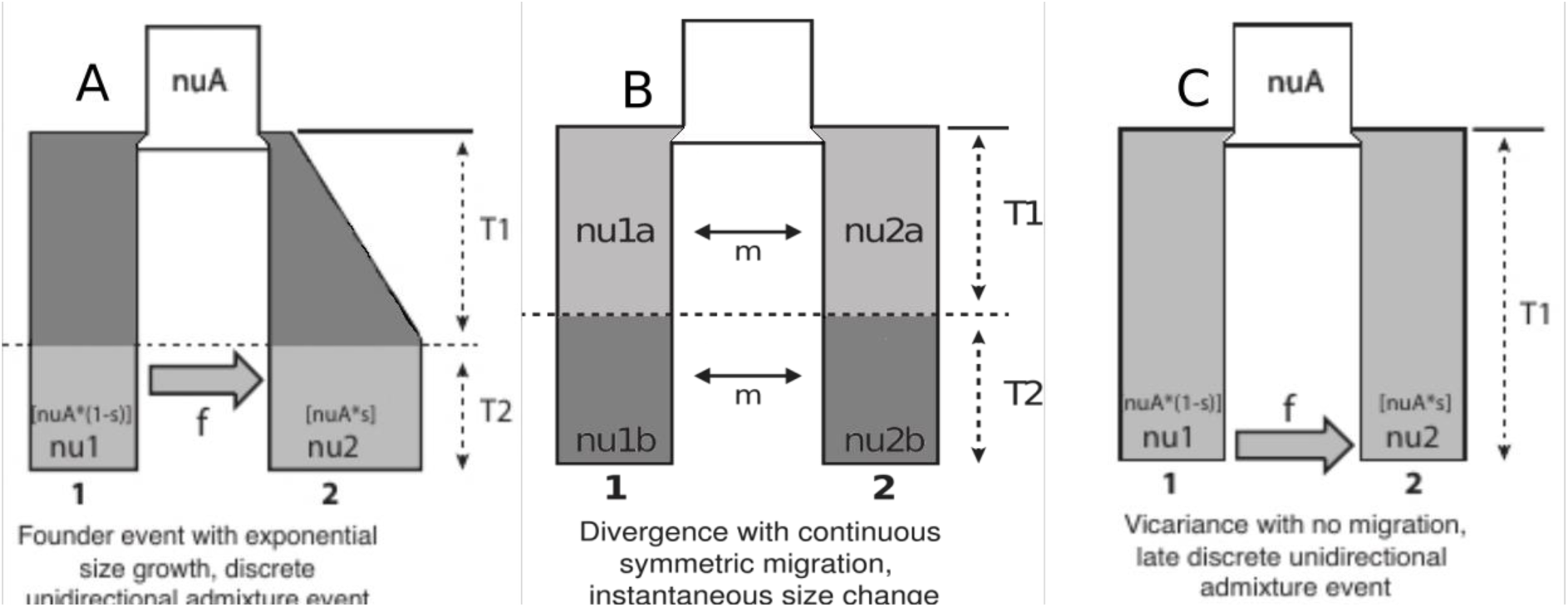
Demographic Model Inference based on the subspecies definition and on the PCA plot. Understanding mechanisms generating parallel genomic divergence patterns among populations is a modern challenge in population ecology, which can widely contribute in the perception of the intraspecific diversification of crop wild relatives. Here we investigated the genomic divergence between three population schemes of *Brassica cretica* using demographic model selection. According to the above results we can support that strict isolation is not recorded between populations. Discrete unidirectional admixture event or continuous symmetric migration was recorded indicating an absence of insuperable barriers in gene flow between populations. Even in the case of taxonomic segregation, where strengthen barriers would be expected, late discrete unidirectional admixture event is corroborated.

### Demographic Model Inference based on the PCA plot

Based on the logPCA results we identified 2 populations. The first comprising three individuals (B, C, D) and the second containing one (A). It is important to note that despite the fact that the A, B, and C plants were sampled from Central Greece and D from Crete, logPCA shows that the Cretan individual is genetically closer to B and C. The distances of A and D to the B-C cluster have small difference, as a result we generated an additional population schema grouping together A, B, C and D as another subpopulation.

For the first grouping, the “Founder event and discrete admixture, two epoch” model, was selected as the most possible demography model (Figure 3B). The second grouping resulted in the “Divergence with continuous symmetric migration and instantaneous size change” as the best model to explain the data (Figure 3C). The first model specifies that the original population split into two subgroups that allowed symmetric migration between them, continuing the population size of each subgroup changed, whereas the second model allows the subpopulations to migrate as the time progresses and the second subpopulation experiences a population size change. The joint 2 population AFS for the real and the simulated data, as well as their difference (residues) are shown in Figure 4.

**Figure 4:**
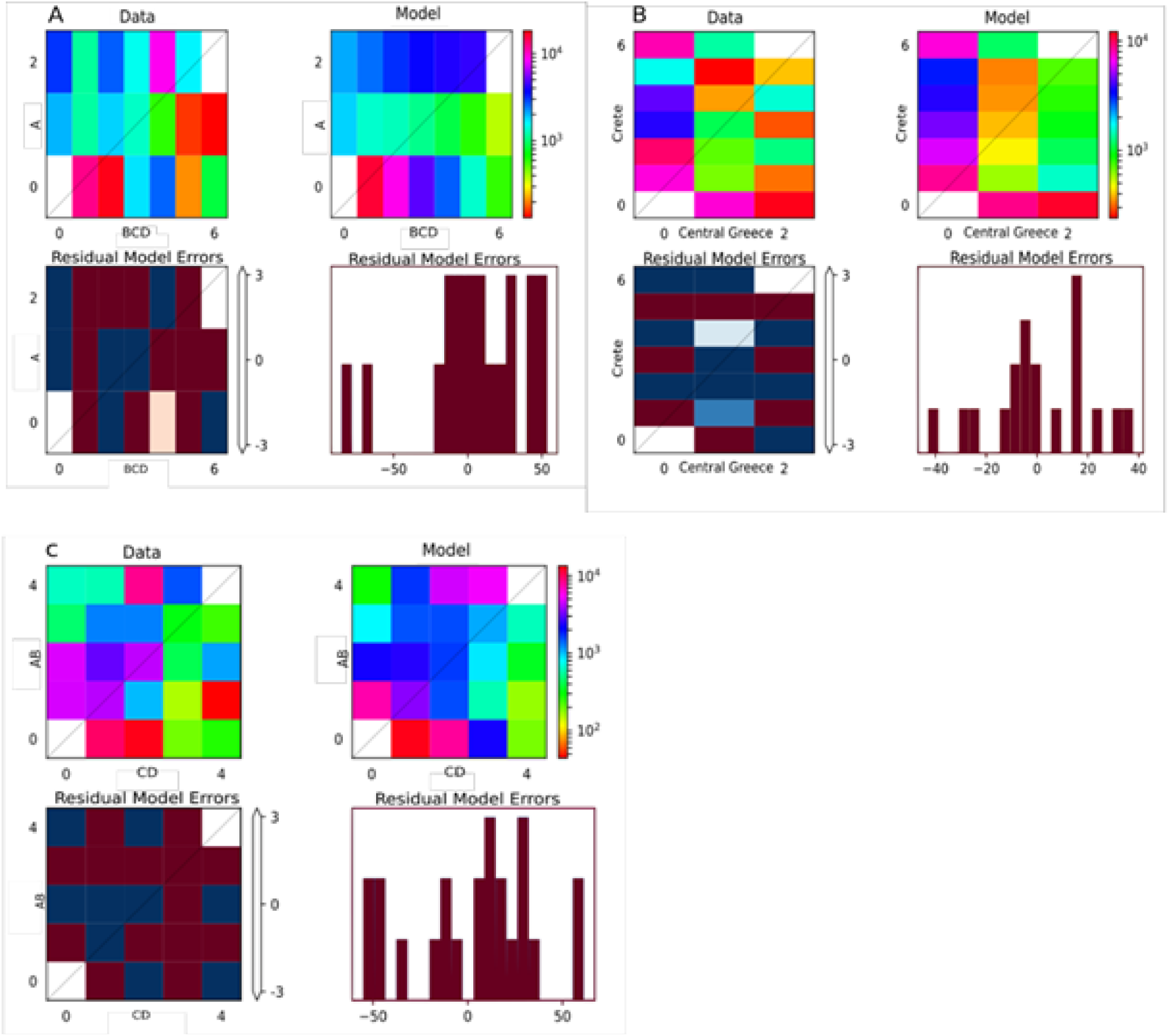
From top right to bottom left: 1) Calculated AFS from *B. cretica* data sets, split by populations. 2) Simulated AFS of the best fitting model from final *dadi* simulations. 3) Heat-map of the residual errors from the comparison between real and simulated AFS. 4) Barplot of the same comparisons.

In all grouping definitions, it is apparent that populations are not isolated. This is considerable gene flow between all possible groupings of the populations. Especially, in the subspecies-based grouping, the inferred model proposes introgression between the two groups, i.e., massive, directional gene flow. Thus, the genetic data suggest that the subspecies separation of *Brassica cretica* plants may be, in fact, not supported by the data. The parameter values for all inferred demographic models as well as the AIC scores of the competing models are presented in the supplementary tables S1, S2 and S3.

The above finding poses the need for further studies concerning the potential gene flow between populations of *B. cretica* and their effects in adaptive traits in both *in situ* and *ex situ* conservation strategies, as well as in cases of genetic improvement especially with newly introduced genes (Snow 2002). There is already evidence of gene flow between wild and crop types of *Brassica* (Prakash *et al.* 2011). Similar concerns have been expressed in the case of crop wild relatives of rice (Thomas *et al.* 2017, Chen *et al.* 2004), which further encourage the incorporation of the followed methodology, that is the demographic model selection in the crop wild relatives research. Of course habitat suitability should also be taken into consideration (Thomas *et al.* 2017, Smykal *et al.* 2017), since ecological factors may also influence the directions and the spatial patterns of gene flow but in the absence of georeferenced data it was necessarily out of scope of the current article. Nevertheless, in future studies a combination of the followed methodology with Ecological Niche Modelling (ENM) (Peterson 2006, Elith & Leathwick 2009) is highly recommended.

In the case of taxonomic segregation, the vicariance-driven divergence with no migration in the early stages indicates that the two taxa typically formed as the result of novel and/or emerging geographical barriers, possibly in combination with genetic drift and/or with the contribution of local adaptation for some traits. Concerning whether non-ecological vs ecological process of genetic isolation took place (Rundle & Nossil 2005), we cannot resort to single explanation since our data are not adequate for such an inference. The late discrete unidirectional admixture event conforms the classical view that in different periods in the evolutionary history of a taxon, different factors (ecological and/or non-ecological) may contribute in the process of speciation inducing or failing to complete it (Nosil *et al.* 2009). Nevertheless, taking into consideration the prevailing hypothesis that plant diversification in the Aegean region is driven by random rather than adaptive differentiation among isolated populations (Strid 1970, Bittkau & Comes 2005, Thompson 2005, Edh *et al.* 2007), we can consider genetic drift as a possible scenario for this population scheme. Apart from that it is worthy of mention that a few studies using population and landscape genetics approaches in *Brassicaceae* have already revealed a significant signal indicating local adaptation (Franzke *et al.* 2011). Smykal *et al.,* (2018) also supported that most of the variation they detected within and between populations of wild pea in northern Fertile Crescent reflects genetic processes such as drift, founder effect and infrequent out-crossing with related individuals, rather than environmental selection pressure.

Unidirectional gene flow has also been reported in cases of other organisms, such as in the case of two lizard subspecies, where gene flow from one subspecies (*Podarcis gaigeae* subsp. *weigandi*) into another (*Podarcis gaigeae* subsp. *gaigeae*) but not in the other direction, recorded by Runemark *et al*,. (2012). In our case, it takes place from the *B. cretica* subsp *nivaea* into the *B. cretica*. Flower colour might be an explanatory factor of the unidirectional admixture event, as in *B. cretica* subsp. *nivea* it is white, while in *B. cretica* may be vary from white to bright yellow but this explanation contradicts with Edh *et al*., (2007), who supported that there is no evidence that flower colour has had in their study any significant effect on gene flow via pollen among the investigated *B. cretica* populations. Nevertheless, in case the view of Edh *et al*., (2007) is depended on the sensitivity of the selected markers (nuclear and chloroplast microsatellites) this flower-coloured based explanation remains standing. Baack *et al.,* (2015) report several cases of pre-pollination reproductive isolation related with flower colour and pollinator behavior.

However, independently whether population genomic divergence is driven by non-ecological or ecological underline mechanisms, the consequences of this late unidirectional admixture event possibly contributed to the high uncertainty or absence of clear consensus of the status of these taxa, which already reported by Edh *et al*., (2007). This is also in line with the treatment of these taxa in the recent Vascular Flora of Greece (Dimopoulos *et al.* 2013), where the taxon *B. cretica* subsp. *nivea* has not been suggested as a standing subspecies.

In the case of non-taxonomic segregations, that is the case of genomic-variation based population schemes, both divergence and founder event were recorded as split mechanisms of the original population, while continuous symmetric migration and discrete unidirectional admixture event in late epoch respectively were specified. In the literature in population genetics, migration and gene flow are often used interchangeably (Tigano & Friesen 2016). Nevertheless, migration refers to the movement and dispersal of individuals or gametes, and gene flow for the movement of alleles, and eventually their establishment, into a genetic pool different from their genetic pool of origin (Endler 1977, Tigano & Friesen 2016). In our case a more appropriate term to use for migration would be dispersal, as migration is mainly used for animals, incorporating also the seasonal movements.

According to the classical view, in the case of founder effect, the genomic variation may be a result in the absence of selection, consequently in this case we can eliminate the role of environment from consideration as an important contribution to genetic variation, while in the case of divergence, the genomic variation may be a result of selective process strengthening the role of environment. Nevertheless, despite predictions on the disruptive effect of gene flow in adaptation, when selection is not strong enough to prevent the loss of locally adapted alleles, an increasing number of studies show that gene flow can promote adaptation, that local adaptations can be maintained despite high gene flow, and that genetic architecture plays a fundamental role in the origin and maintenance of local adaptation with gene flow (Tigano & Friesen 2016). Thus, in the genomic era it is important to link the selected demographic models with the underlying processes of genomic variation because, if this variation is largely selectively neutral, we cannot assume that a diverse population of crop wild relatives will necessarily exhibit the wide ranging adaptive diversity required for further crop improvement.

## Supporting information

Supplementary Table S4: xls file containing Information for all genomes obtained from the GenBank

## Conflict of Interest Statement

The authors declare that they have no conflict of interests.

## Authors’ contributions

P.F.S. and D.J.S. designed the research. A.K., V.A.M., L.B. and P.P. performed the research. S.P., D.J.S., P.P., A.K., and P.F.S. analyzed the data. P.P., A.K., S.P., D.J.S. and P.F.S. wrote the paper.

## Acknowledgments

P.F.S and D.J.S. were supported by a grant from the Gatsby Charitable Foundation. The authors would like to acknowledge Dr Karen Moore and the Exeter Sequencing Service at University of Exeter, for technical assistance with DNA genome sequencing.

## Supplementary Material

**Supplementary Table S1A:** Top 3 AIC relative weights models with BCD-A cluster:

| Model Name | Relative AIC Weight |
| --- | --- |
| Founder event and discrete admixture | ~1 |
| Divergence with ancient symmetrical migration | 1.67e-91 |
| Divergence with no migration | 5.08e-211 |

**Supplementary Table S1B:**
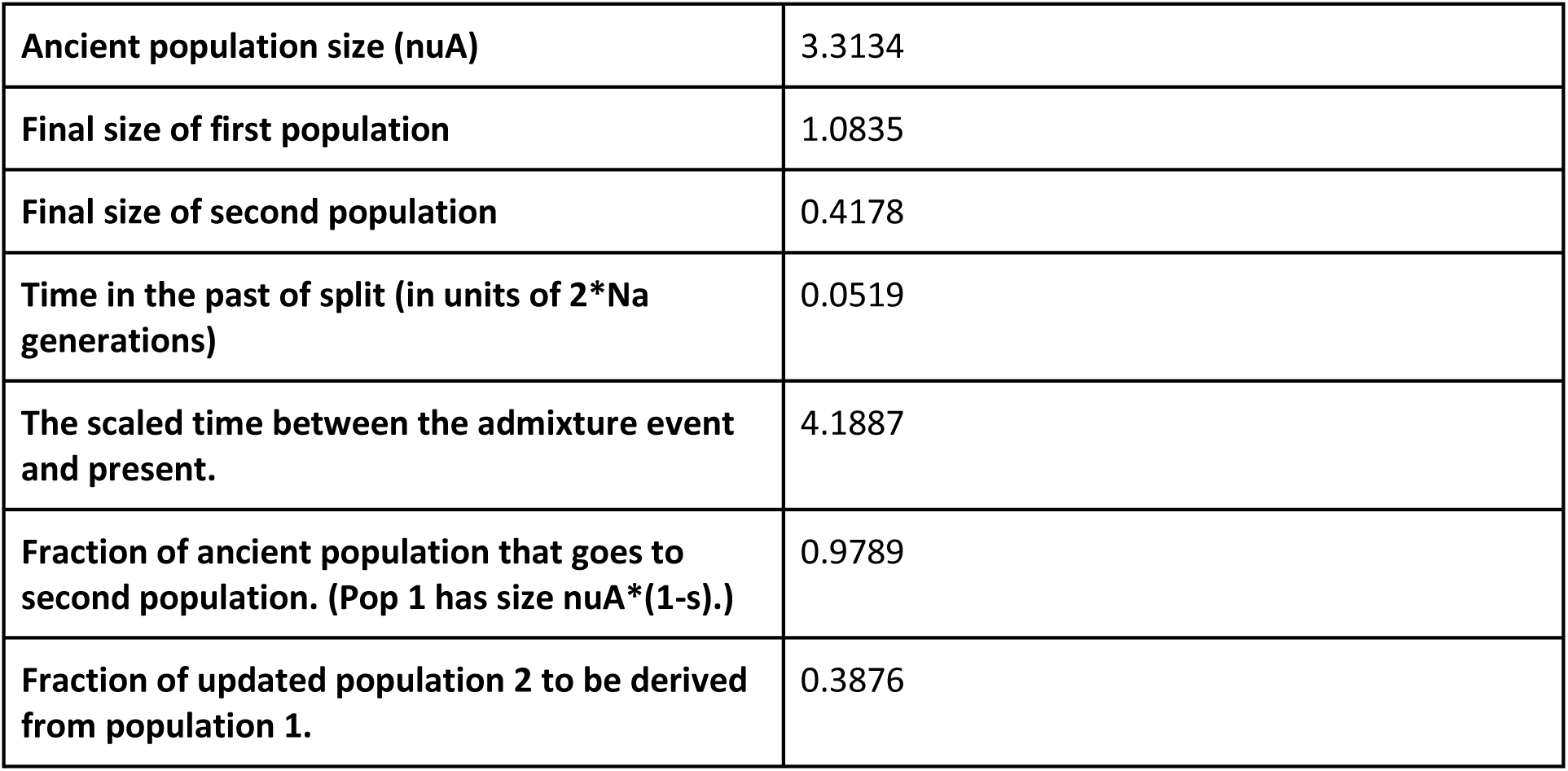
Parameters of optimal model on BCD-A cluster:

**Supplementary Table S2A:**
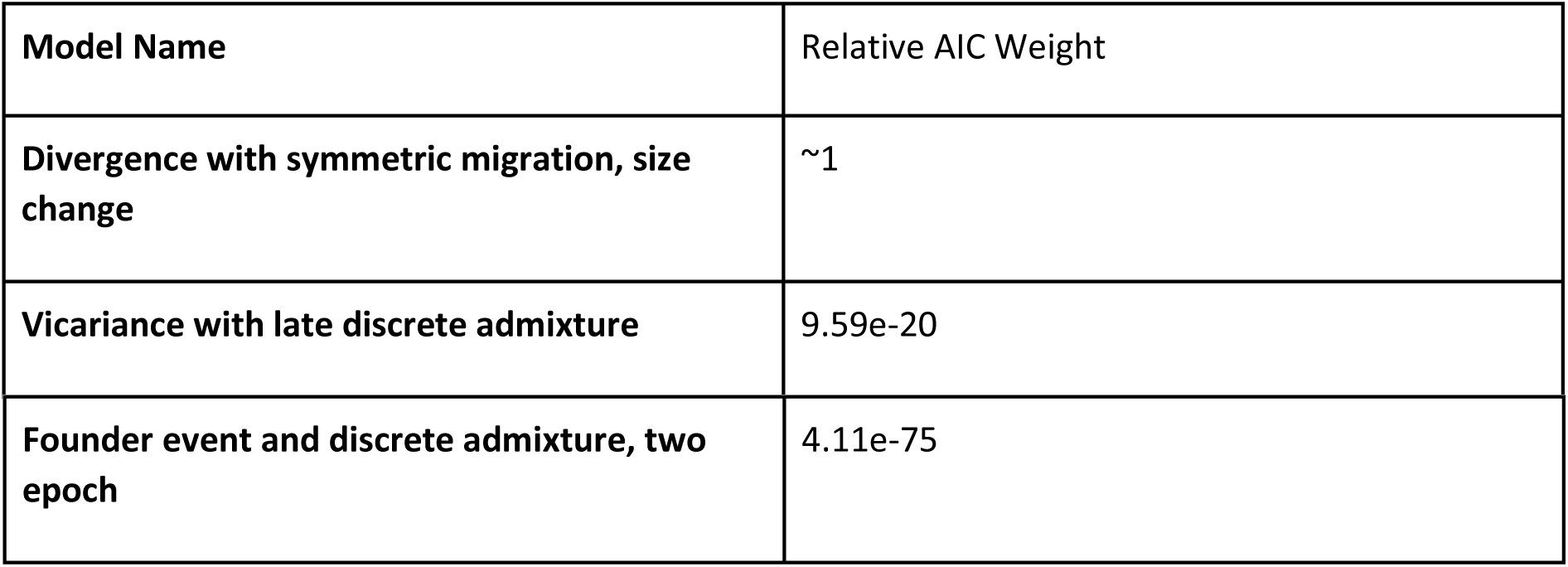
Top 3 AIC relative weights models with ABC-D clusters:

**Supplementary Table S2B:**
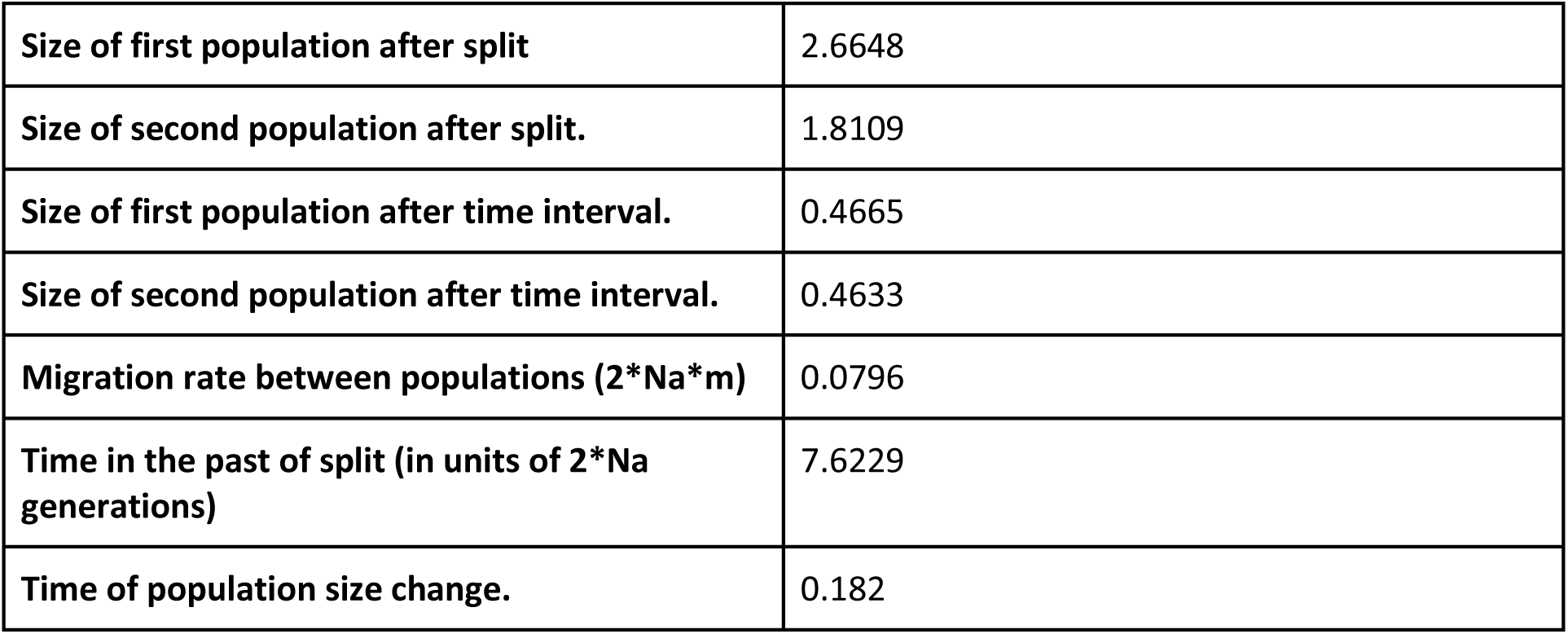
Parameters of optimal model on ABC-D cluster:

**Supplementary Table S3A:**
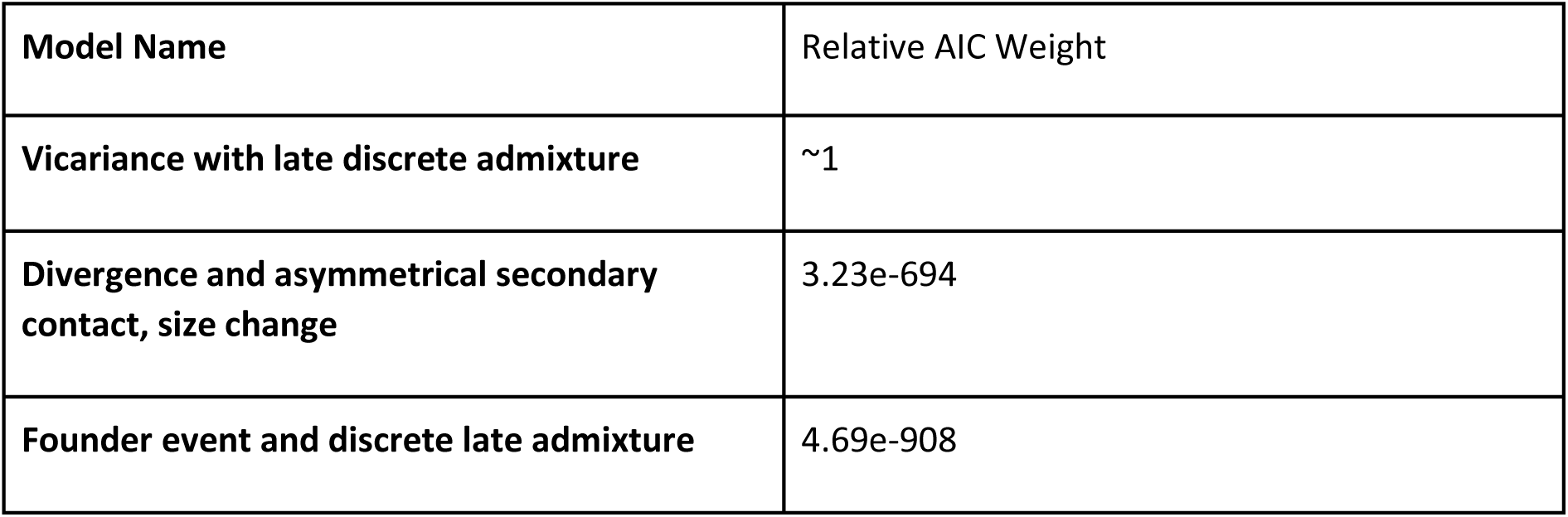
Top 3 AIC relative weights models with the AB-CD clusters:

**Supplementary Table S3B:**
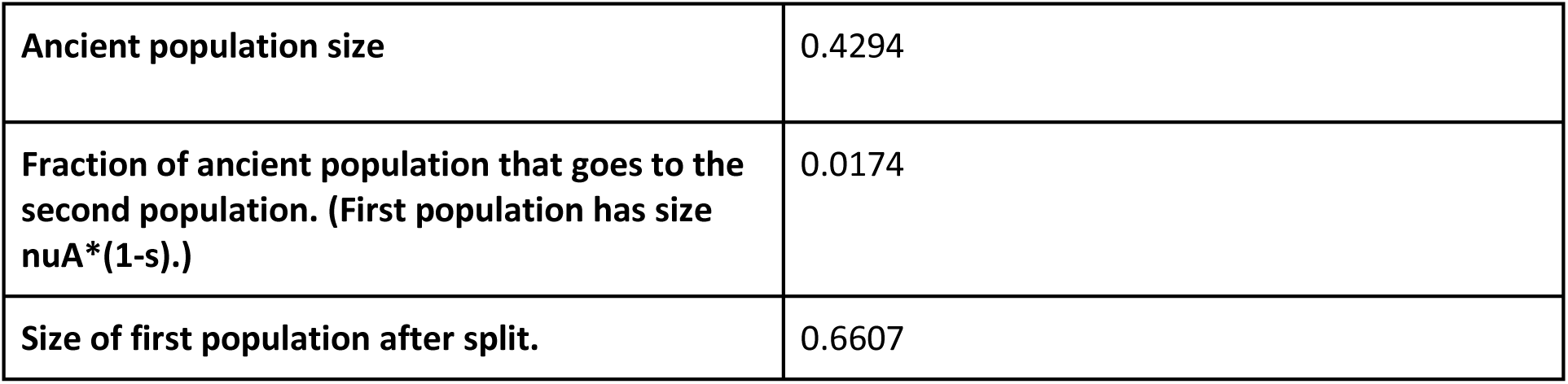
Parameters of optimal model on AB-CD clusters:

**Supplementary Table S4:**
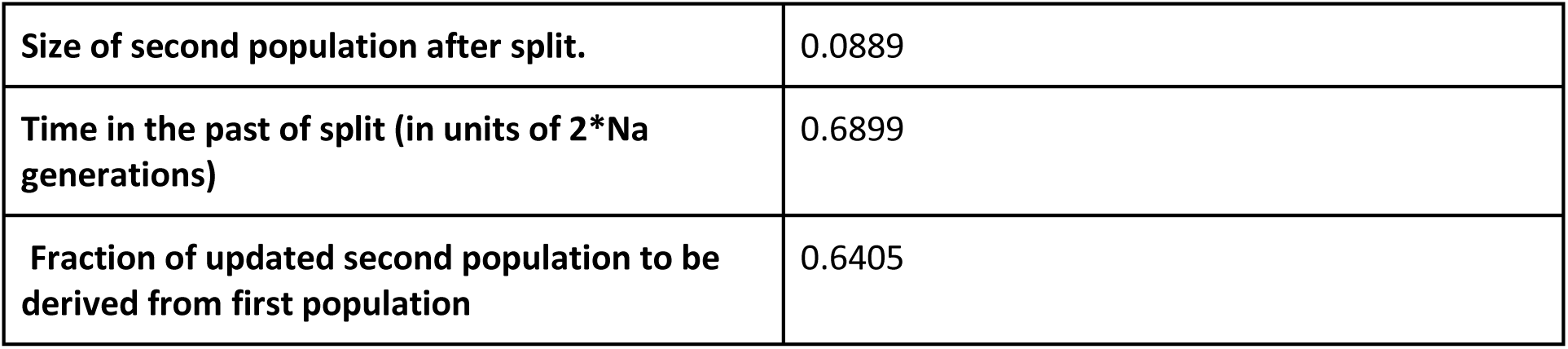
*csv file* containing Information for all genomes obtained from the GenBank.

